# Long-read sequencing of SARS-CoV-2 reveals novel transcripts and a diverse complex transcriptome landscape

**DOI:** 10.1101/2021.03.05.434150

**Authors:** Jennifer Li-Pook-Than, Selene Banuelos, Alexander Honkala, Malaya K. Sahoo, Benjamin A. Pinsky, Michael P. Snyder

## Abstract

Severe Acute Respiratory Syndrome Coronavirus 2, SARS-CoV-2 (COVID-19), is a positive single-stranded RNA virus with a 30 kb genome that is responsible for the current pandemic. To date, the genomes of global COVID-19 variants have been primarily characterized via short-read sequencing methods. Here, we devised a long-read RNA (IsoSeq) sequencing approach to characterize the COVID-19 transcript landscape and expression of its ∼27 coding regions. Our analysis identified novel COVID-19 transcripts including a) a short ∼65-70 nt 5’-UTR fused to various downstream ORFs encoding accessory proteins such as the envelope, ORF 8, and ORF 9 (nucleocapsid) proteins, that are relatively highly expressed, b) novel SNVs that are differentially expressed, whereby a subset are suggestive of partial RNA editing events, and c) SNVs at functional sites, whereby at least one is associated with a differentially expressed spike protein isoform. These previously uncharacterized COVID-19 isoforms, expressed genes, and gene variants were corroborated using ddPCR. Understanding this transcriptional complexity may help provide insight into the biology and pathogenicity of SARS-CoV-2 compared to other coronaviruses.

## Introduction

The Severe Acute Respiratory Syndrome Coronavirus 2, SARS-CoV-2, is a highly infectious betacoronavirus and the cause of the coronavirus disease 2019 (COVID-19) responsible for the current 2020 global pandemic (as of December 2020; over 88 M cases worldwide based on Worldometer.info). The disease predominantly affects the lungs and causes varying degrees of severity and symptoms that range from asymptomatic to mild cases of fatigue, fever/chills, cough, ache, and loss of smell, although up to 20% of cases will be severe with pneumonia, dyspnea, and acute respiratory distress syndrome^1,2^. It has caused more infections to date than other severe coronaviruses, such as SARS-CoV-1 and Middle East Respiratory Syndrome Coronavirus (MERS-CoV), combined^3^. The pathway of infection for both SARS-CoV-1 and 2 is via a receptor binding domain (RBD) on the C-terminal of the spike protein S1 subunit protruding from the viral capsid that binds to human angiotensin-converting enzyme 2 (hACE2), which is present at high levels in pulmonary epithelia. Antibodies in convalescent patients are primarily targeted to this RBD region and accordingly vaccine development has been focused on this domain^4^. Notably, there are four distinct amino acid variants (G, V, E, G) present in the RBD domain of SARS-CoV-2 that increase the affinity of SARS-CoV-2 binding to hACE2 relative to other coronaviruses^5,6^. Developing a better understanding of the sequence variation and a comprehensive transcript map of SARS-CoV-2 across global populations is important to better understand viral biology, evolution, and the development of improved therapies against COVID-19 infection.

In early January 2020, the first near complete SARS-CoV-2 genome was released to the GISAID database and is widely used as the primary COVID-19 reference genome (NC_045512). As of December 2020, over 47,600 SARS-CoV-2 sequences have now been recorded in the NCBI database. SARS-CoV-2 is a large RNA virus with a ∼29.8kb genome sharing up to 79% sequence identity with SARS-CoV-1; other family members include HCoV-229E, HCoV-OC43, HCoV-NL63, and HCoV-HKU1, which cause common colds^3,7^. SARS-CoV-2 contains 27 proteins generated by 14 open reading frames (ORFs) (Fig. 1a). The long ORF1a and ORF1b regions are translated as one large peptide and then post-translationally cleaved into 15 nonstructural proteins (nsp), including nsp 1-10, 12a/b, 13-16, an RNA dependent RNA polymerase (RdRP), helicase (H), exonuclease (Ex) and endonuclease (En)^8^. The nsp proteins include two viral proteases, nsp 3 (papain-like protease) and nsp 5 (3C-like protease), involved in the post-translational processing of ORF1a/b. Further downstream are the structural proteins, including the two subunits of the spike surface protein, S1 and S2, which contain the least-conserved regions among coronaviruses, as well as the regions encoding the receptor binding motif (RBM) within the RBD and a polybasic cleavage site between S1 and S2^9^. Following the S2 region are seven accessory proteins interspersed between the envelope protein (E), membrane glycoprotein (M) and nucleocapsid phosphoprotein (N) regions. The most common methods for the molecular detection of COVID-19 involve extraction of nucleic acids from nasopharyngeal swabs, conversion of RNA to cDNA, and amplification using qPCR probes designed to conserved sites within the SARS-CoV-2 genome, including regions in the E-protein and RdRP genes.

**Fig. 1:**
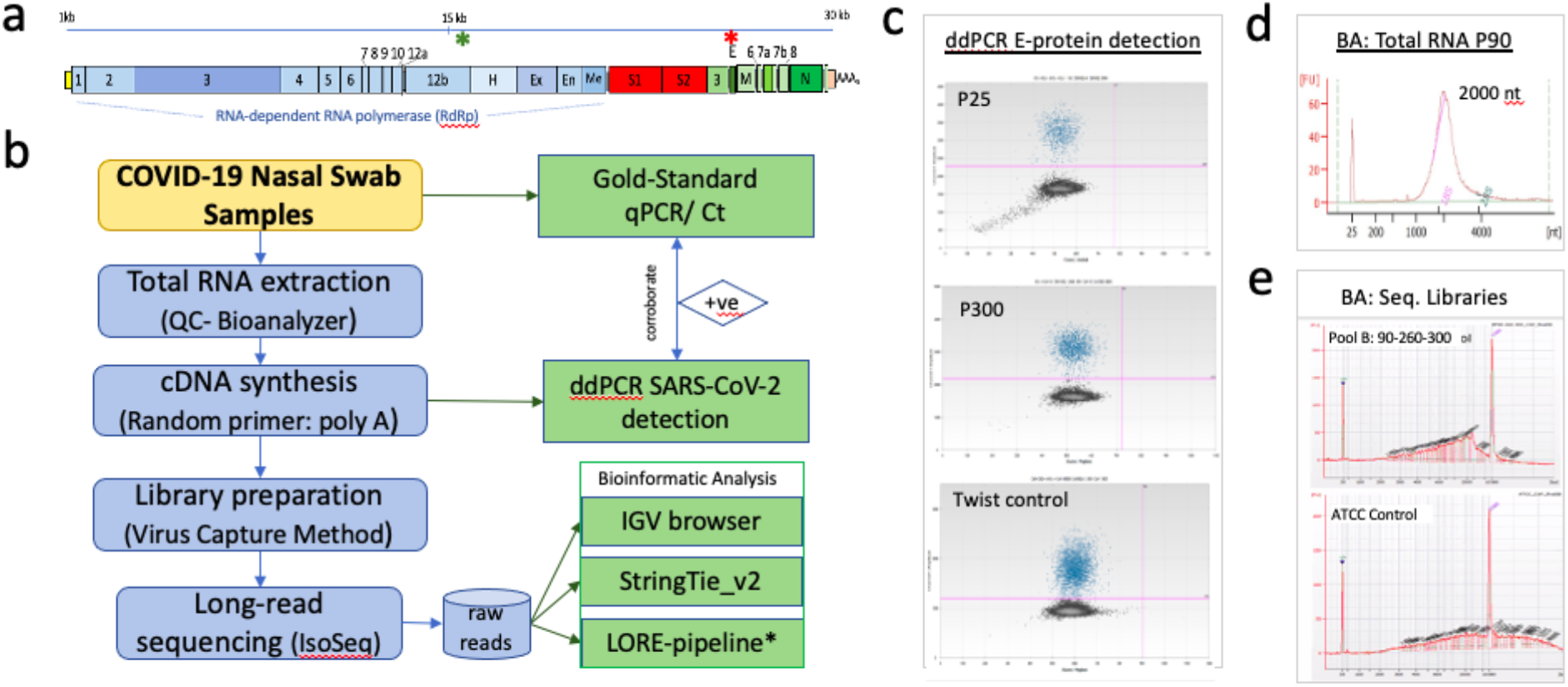
Characterization of RNAs from COVID-19 positive samples. (a) Schematic of SARS-CoV-2 gene structure based on reference GenBank: MT008022.1 denoting nonstructural proteins (nsp 1-16 in blue) including RNA dependent RNA polymerase (RdRp), Helicase (H), Exonuclease (Ex) and Endonuclease (En), the spike protein subunit 1 and 2 (in red), and structural proteins in green (ORF 3-10, whereby ORF 4 is the Envelope protein (E), ORF 5 is the Membrane protein (M) and ORF 9 is the Nucleocapsid protein (N). The red asterisk shows the location of primers/probes used for Gold-Standard qPCR testing of this virus (E-protein), and the green asterisk shows the location of a second probe set designed RdRp of the SARS-CoV-2 virus. (b) Flowchart of long-read sequencing methodology. The IGV browser and StringTie_v2 was used for the long-read analysis. The <u>L</u>ong-<u>Re</u>ads-pipeline (LORE) was developed to further analyze gene rearrangements for this work. (c) Confirmation of COVID-19 positive samples using parallel Gold-Standard E-protein primer/probes via ddPCR with positivity in the FAM channel (blue dots). Panel: Patient samples P25 (top), p300 (middle) and Twist synthesized SARS-CoV-2 control (bottom). (d) Representative Sample Bioanalyzer (BA) data of total RNA extracted from nasopharyngeal swabs from a patient 90 (p90) testing positive for COVID-19. RNA transcripts range in size ∼200-6000 nts (y-axis). (e) Bioanalyzer (BA) QC of final capture of SMRTBELL long-read sequencing libraries (Top Panel: Patient Pool B Samples P90-P260-P300, Bottom Panel: ATCC SARS-CoV-2 control).

The mechanism of gene expression in positive strand RNA viruses like SARS-CoV-2 is complex with three hypothesized mechanisms for the production of mature RNAs, subgenomic (SG) RNAs, from a large negative strand (-ve) RNA template. These proposed mechanisms include using 1) the -ve strand internal promoter for transcription initiation; 2) prematurely terminated -ve strands acting as heterogeneous templates for production of each SG RNA species; or 3) discontinuous RNA synthesis of the -ve strand template^10^. The vast majority of COVID-19 genomes and transcripts sequenced to date have used short-read methods which, although powerful for rapid sequencing results and sequence comparison, nonetheless yield limited data on the different SARS-CoV-2 transcripts. In particular, we do not know the catalog of the transcripts produced by the virus and the levels of these transcripts, nor do we know which single nucleotide variants are associated with the different isoforms and mechanisms by which the variants might have formed. Such information will be valuable for understanding the basic biology of this virus and its pathogenicity^11^, particularly in the context of emerging new strains, such as the recently identified B7.1.1 variant that may have increased transmission characteristics^12,13^.

To better understand the SARS-CoV-2 RNA landscape, we adapted a long-read PacBio sequencing technology to characterize SARS-CoV-2 transcriptomes from multiple COVID-19 patients. We developed an analysis pipeline to describe a suite of COVID-19 transcripts, several of which appear to be full-length, and describe their levels, a subset validated by ddPCR. We discovered novel 5’- and 3’-end UTRs and unusual associated transcriptional rearrangements. These UTRs often included novel repetitive sequences. We further associate a natural single nucleotide variant (SNV) associated with specific spike protein isoforms with an expression decrease relative to the “wildtype” nucleotide. Taken together, these data provide new insight into the overall transcript-processing landscape of SARS-CoV-2, including partially RNA edited transcripts, and provide an improved framework for understanding COVID-19 variants, gene isoforms, and their expression mechanisms.

## RESULTS

### Characterization and quality of SARS-CoV-2 RNA

We adapted a method for long read sequencing of SARS-CoV-2 transcripts using Pacific Biosciences SMRT Sequencing technology (Fig. 1). Patients from the San Francisco Bay Area were tested for COVID-19 using nasopharyngeal swabs and an FDA-Emergency Use Authorized realtime RT-PCR method, with Cycle threshold (Ct) values recorded as a proxy for abundance (Supplementary Fig. 1). We independently confirmed the presence and level of SARS-CoV-2 sequences using ddPCR (Fig. 1c). This is using the parallel E-protein primer/probes of the Gold-Standard SARS-CoV-2 qPCR detection method, but with the additional dual-channel fluorometric capacity of the ddPCR platform (Materials and Methods). From a set of 24 COVID-19 positive samples, six of high quality were selected for sequencing and analysis (Fig. 1b). Total RNA isolated from these high-quality nasopharyngeal samples showed an average transcript size of 1500-3000 nt using the Bioanalyzer (Fig. 1d) as compared to other samples that have many lower sized RNAs, suggestive of some degradation (Supplementary Fig. 1a). The nasopharyngeal RNAs of these six samples primarily produced a single band peak at 2 kb ranging from ∼1.5 to 4 kb while unexpectedly rRNAs from the human host were not detected in the Bioanalyzer data. Two SARS-CoV-2 RNA controls were also analyzed in parallel with the patient samples, 1) a Twist Bioscience SARS-CoV-2 (MT199235 – USA/CA9/2020) synthetic control containing six non-overlapping 5 kb fragments and 2) SARS-CoV-2 RNAs isolated from Vero E6 mammalian cells transfected with a 30 kb cloned isolate (MN985325.1 −2019-nCoV/USA-WA1/2020) supplied by ATCC (ATCC® VR-1986D™). These controls also produced a single band using the bioanalyzer (Supplementary Fig. 2). Overall, we conclude that SARS-CoV-2 RNAs of 1.5 to 4 kb in size dominate the pool of nasopharyngeal RNAs in infected patients.

### Long Read sequencing of SARS-CoV-2

Because patient SARS-CoV-2 RNAs are low in abundance, to optimize concentrations and potentially identify as many SARS-CoV-2 transcripts as possible, two sets of total RNA were pooled from three different high-quality COVID-19+ samples: Pool A (Patients 25, 125, 290) and Pool B (Patients 90, 260, 300). Universal Human Reference RNA (UHRR, Agilent) was included to balance the RNA concentrations in the pools. cDNA was generated from total RNA using a 9:1 ratio of random primers:anchored oligo-dT primer and used to generate cDNA libraries ranging from 1500-3000 bp. SARS-CoV-2 viral transcripts were captured using custom designed 120bp IDT probes. Sequencing was done on a single SMRT Cell on the PacBio Sequel system. This resulted in 59,674 circular consensus sequence (CCS) reads from Pool A and 42,629 from Pool B. Since the UHRR was spiked into the sample to increase RNA concentrations, the majority of sequenced reads matched the UHRR controls (and not human host sequence as observed by lack of rRNA), corroborating the observation that prior to spike-in, most real-world sample reads are viral (Table 1). The average length of the reads was ∼1915 nt for Pool A and ∼1871 nt and in Pool B (Table 1), and range in size from 300 nt – 5 kb (Supplementary Fig. 2a-b).

**Table 1:** Coverage and length of long-reads CCS (PacBio)

|  | BP25-120-290<br>(Patient) | BP90-260-300<br>(Patient) | Twist (+ve<br>control) | ATCC (+ve<br>control) |
| --- | --- | --- | --- | --- |
| #CCS reads<br>(3pass) | 59,674 | 42,629 | 67,705 | 24,684 |
| #mapped CCS | 7,365 | 9,709 | 26,581 | 5,278 |
| % mapped CCS | 12.34% | 22.77% | 39.26% | 21.38% |
| Mean mapped<br>CCS RL | 1871bp | 1915bp | 1757bp | 1785bp |
| Max mapped CCS<br>RL | 5191bp | 4022bp | 5045bp | 5110bp |

### Diversity in transcript isoforms: Known and novel transcripts

To capture the variety of SARS-CoV-2 RNA transcripts and their expression levels from the long-reads, we used a combination of StringTie v2.1.3 as well as our own custom pipeline to assemble reads into transcript classes (LORE pipeline, Supplementary Fig. 3). The two pooled sets of data from the six COVID-19 positive samples revealed at least fifteen distinct isoforms of SARS-CoV-2 transcripts based on the presence of presumed fusion events and that many of these contained polyA tails (although this does not include the variable 5’ ends, which are heterogeneous). Interestingly, only two of these isoforms were shared among the pooled sets, Ai2/Bi2 and Ai7/Bi3 (Fig. 2). In addition, three sets of sequences detected, Ai7, Bi1 and Bi3 at relatively low abundance, lacked spliced regions or polyA tails and may thus represent either subgenomic fragments or parts of transcripts. Finally, several fusion transcripts were found in very low abundance <5 and are described further below. In aggregate, the transcripts fully covered the reference genome isolate Wuhan-Hu-1 (MT008022, NC_045512.2).

**Fig. 2:**
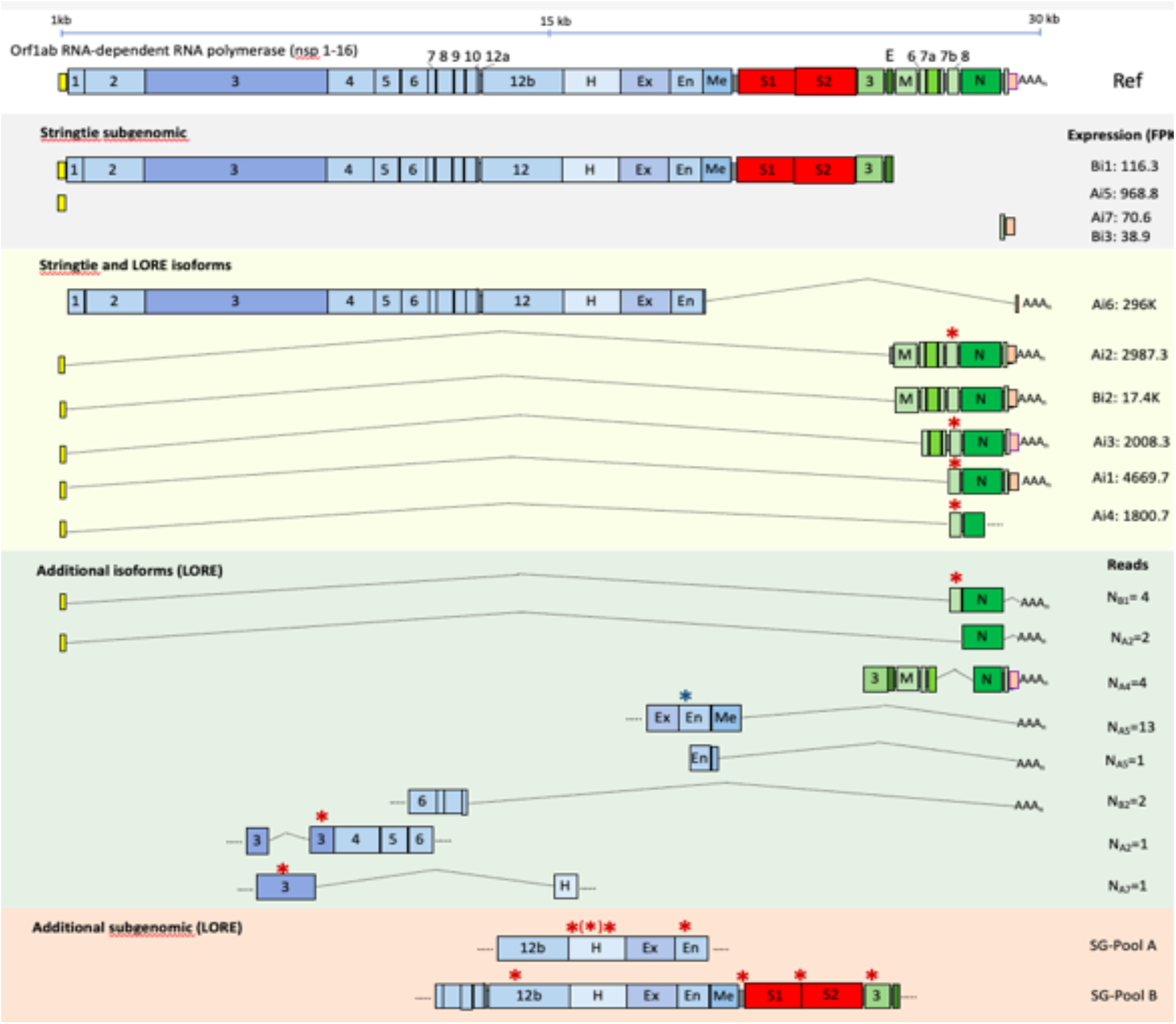
Schematic representation of the SARS-CoV-2 genome, isoform landscape and expression levels. Colored boxes represent the ORFs based on Wuhan-Hu-1 (MT008022, NC_045512.2) as the reference genome. Yellow, blue, red, green, and orange boxes are 5’-UTR, non-structured proteins (nsp), spike protein (S1 and S2), seven accessory proteins and 3-UTR. Helicase (H), exonuclease (Ex), endonuclease (En) and methyltransferase (Me) are among nsp proteins (blue). Envelope (E), membrane protein (M) and nucleocapsid (N) are the identified structural proteins (green). Isoforms and expression levels (in FPKM) identified through the StringTie_v2 pipeline are denoted as Ai1-7 and Bi1-3 from Pools A and B, respectively. The number of unusual isoforms detected through the developed LORE pipeline are denoted as N_A(1-7)_ and N_B(1-2)_ from Pools A and B, respectively. Red asterisks show SNV location, blue asterisk is wildtype nucleotide within an isoform in Pool A. Asterisk in a bracket (*) represent a subpopulation of SNV among the reads that is the wildtype nucleotide, suggesting partial editing.

The majority of transcripts identified ranged in size from 300 nt to ∼5kb and their relative abundance based on read frequency (which may be biased) is shown in Fig. 2. We believe that many of these are full length transcripts as they contain 5’ genomic regions as well as poly A tails. Indeed, on average, 13.5% and 6.0% of transcripts in Pools A and Pool B, respectively, contained poly A tails ranging from 15-40 A’s in length, which were found at the end of the majority of identified isoforms. The majority of polyA transcripts were found on the transcript of the accessory proteins isoforms, particularly the penultimate nucleocapsid (N) ORF (Fig. 2 green). The high prevalence of this polyA transcript may be a result of 3’ bias of the RT-PCR method used to generate the sequencing libraries.

### Expression Levels of SARS-CoV2 ORFs and transcripts

All 27 ORFs were expressed, albeit at varying levels, in the nasopharyngeal samples (Fig. 2). The majority of nsp (blue, Fig. 2) were found to be expressed on the same transcript, although at low expression levels relative to the accessory proteins (in Pool A), whereas the spike protein had expression levels below the limit of detection via StringTie in Pool A. However, the spike protein was detectable as part of a long isoform also containing nsp and partial accessory protein (green, Fig. 2) in Pool B. Each SARS-CoV-2 gene was also observed for the ATCC controls, albeit at more homogenous expression levels, as expected for cloned products.

The two SARS-CoV-2 RNA controls (Twist and ATCC controls) were also sequenced in parallel with patient samples. As expected, when mapped to genomic regions, the reads from the synthetic interspaced 5 kb fragments of the Twist control produce five clusters of reads whereas the ATCC control, generated in vitro, has a more distributed expression pattern similar to the COVID-19 patient samples (Fig. 3).

**Fig. 3:**
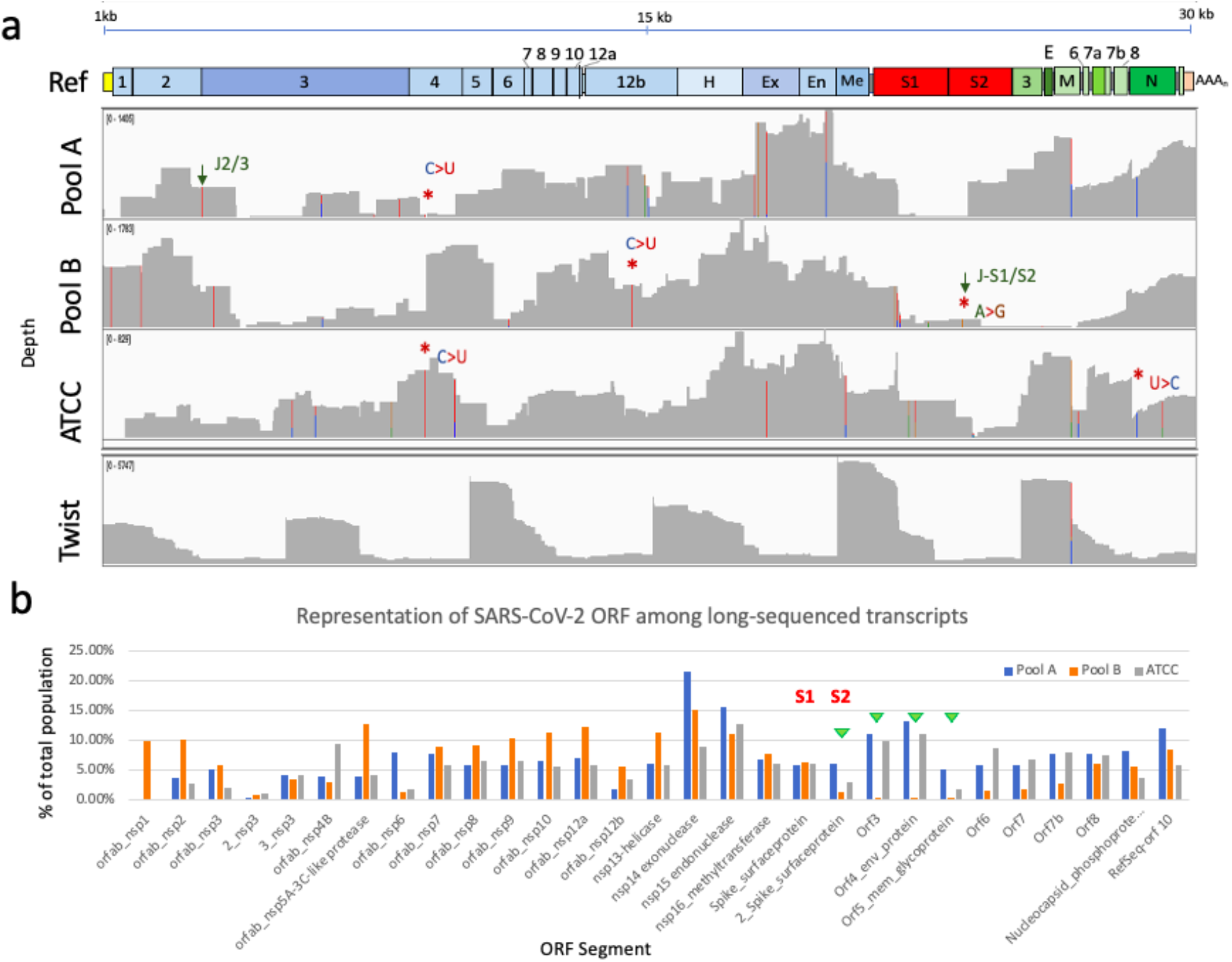
Transcriptome landscape and variants of SARS-CoV-2. (a) IGV Browser (v.2.8.0) depicts long-read distribution and single-nucleotide variants of the two pools (A and B) of COVID-19 patient samples and the two SARS-CoV-2 RNA synthesized controls: ATCC (MN985325, Washington Source) and Twist (MT199235, California Source). Green arrows marked as J2/3 and J-S1/S2 denote a SNV within 10 nucleotides of the junction sites nsp 2/nsp 3, and spike protein subunit 1 and subunit 2, respectively. Asterisk denotes SNV validated through ddPCR (Fig. 4). (b) Bar graph representation of SARS-CoV-2 ORF/gene expression relative to total reads among the long-sequenced transcript population for Pool A (blue), Pool B (orange) and ATCC SARS-CoV-2 control (gray). Note the lower expression in genes downstream of S2 for Pool B samples suggesting an association with the SNV at the spike subnunits 1 and 2 junction (J-S1/S2) depicted in (a).

In the real-world patient samples, up to fourteen distinct isoforms were identified in the population of RNAs, with the most abundant expression (highest FPKM) including four variations of genes at the 3’ end ORFs (Fig. 2, left). These ORFs include part of the envelope protein (E), the membrane glycoprotein (M), ORFs 6-8, ORF 9/nucleocapsid phosphoprotein (N) and ORF 10. Corroborating the fusion hypothesis in coronavirus transcription, a shorter 1-65 nt segment of 5’ UTR, also described by *Kim et al*., was found to be joined or ‘alternatively spliced’ to four varieties of isoforms, including M, N, and ORFs 6-8 (downstream accessory proteins), instead of the annotated full-length 1-260 nt 5’ UTR^10,15^ (Fig. 2-green section). The ATCC control (derived from clones) also showed a variety of full-length isoforms, some altogether lacking the expected full-length 5’ UTR. Segregated isoforms of 5’ and 3’ UTR transcripts were also identified in the population, as a subset of fused isoforms of N protein to poly A, missing ORF 10 and 3’ UTR (N_A1_ and N_B1_). Although the RNAs were heated to denature RNA structures (Methods), both 5’ and 3’ UTRs of coronaviruses are known to have significant secondary hairpin structures^16^ and the reverse-transcriptase methods used to generate the sequencing libraries may be unable to capture the entire 5’ region.

### Discovery of novel single nucleotide variants and their differential expression

Distinct from traditional long-sequencing pipelines^17,18^, the <u>Lo</u>ng-<u>Re</u>ads Analysis (LORE) was developed to additionally identify strings of consecutive genes, non-consecutive regions, as well as characterize the single nucleotide variants (SNVs) transcripts alongside their relative expression levels. A number of expressed novel single nucleotide variants (SNVs) were also discovered in the long read regions as shown in Fig. 3; some of these likely have functional consequences. In order to identify SNVs of high confidence, variant correction based on sequencing depth was applied to correct for platform errors, which include RT-PCR amplified mis-incorporated single nucleotides, as well as the relatively high error rate of indels in raw PacBio long-sequence reads (data not shown). Similarly, the ATCC controls are from cultured Vero cells, potentially introducing variant amplification errors distinct from the original strain. Nine and seven high confidence SNVs were identified in samples from Pools A and B, respectively, with a minimum coverage depth of 25 reads (located >10 nt from the read ends). These SNVs are shown relative to the Twist and ATCC controls in Fig. 3a. Four SNVs overlapped between Pool A and the ATCC control (reference is a Washington source patient). Over 80% of the novel SNVs are C-to-U (9) or A-to-G (4), suggesting an RNA editing (deaminase-like) modification (Supplementary Table 2). Positions 17747, 17858 and 18060 show expression of their alternate variants at 3%, 2% and 4%, respectively, and suggest these are candidate transcripts for partial editing (Figure 4a, purple triangles). Positions 17747 and 18060 alter the coding amino acid, P-to-L and Y-to-C, respectively. Note that the three patient samples P25, P125 and P290 are represented at ratios greater than 20% in Pool A at 0.48, 0.32, and 0.21, respectively (Materials and Methods), and partially edited nucleotides could be representing a temporal event in any (or all) of the patients.

**Fig. 4:**
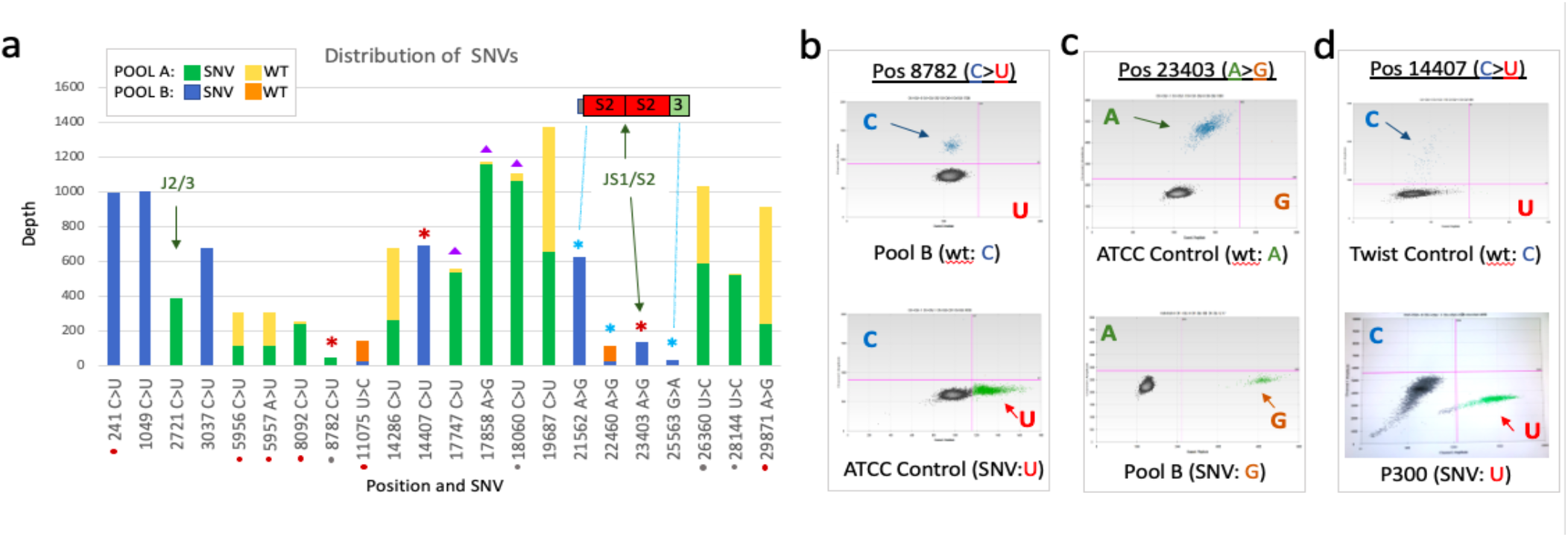
Distribution of single nucleotide variants (SNVs) in COVID-19 patients from the California Bay Area. (a) Bar graph representation of SNV position and depth in Pool A (green/yellow) and Pool B (blue/orange), with a total of sixteen, nine and seven high confidence SNVs, respectively. Red asterisks denote the SNVs corroborated by digital droplet PCR (b-d). Purple triangles are candidates for partial-editing (coverage >500 reads (min. 25 reads), with the wildtype nucleotide expressed at 2-5%). The x-axis is the position and type of SNV, where grey dots denote shared SNVs with ATCC synthetic RNA control. The red dots represent six lower quality SNV candidates, found in A-rich or T-rich regions, including the last nucleotide before poly A tail (position 29871), suggesting a sequencing mis-incorporation. The y-axis is the read coverage depth of the SNV (minimum of 20 reads, non-unique with cut off 3 unique). (b) ddPCR method showed that Pool B patient samples (top panel) contained “C” wildtype nucleotide at position 8782 (within the nsp 4 gene), while the ATCC SARS-CoV-2 control (and Pool A) contained the “U” variant (bottom panel); (c) This is contrasted with ddPCR whereby the ATCC control contained the wildtype “A” variant (top panel) while the Pool B patient samples contained the “G” variant at position 23403 (spike/surface glycoprotein gene). Additionally, a decrease in droplet counts also corroborated the decrease in expression observed in Pool B relative to the ATCC SARS-CoV-2 RNA control profile (Fig. 3). (d) ddPCR also corroborates the wildtype “C” at position 14407 (RdRp) in the Twist RNA synthetic control (top panel), while patient 300 (P300) showed the “U” variant. P300 was one of the samples in Pool B. See Supplementary Table 1 for primer/probe detail. Reference RNA genome GenBank: MT008022.1 (Wuhan) was used. ddPCR allows for dual-channel probes (FAM and HEX) which were designed to specify each nucleotide (wildtype and variant, respectively). FAM (y-axis) is represented by blue dots and HEX (x-axis) is represented by green dots (droplets) in panels b-d.

Other observations include the identification of SNV-type isoforms (Fig. 2, asterisks), as A_i2_ and B_i2_ which are similar in shortened 5’UTR fusion with accessory ORFs but differ in a single SNV at position 28144; and have >5-fold difference in expression levels. Among Pool A transcripts, N_A5_ isoform of nsp 14-16 (Ex, En, Me) fused to poly A, contain the wildtype “U” at position 26360, while a subset of tiled reads (concatenated SNVs found in nsp 12b, 13-15), categorized here as subgenomic (Fig. 2; SG-Pool A asterisks), contain the “C” SNV at the respective position. Interestingly, two SNVs were found near (1-2 nt) protease cleavage site junctions that may have important functional consequences. A C-to-U change was found 1 nt from the nsp2/3 cleavage site at position 2721, and a G-to-U change was identified 1 nt from the spike subunit 1 and subunit 2 cleavage site at position 23403 (Fig. 3a and 4a, green arrows). Interestingly, this latter site is associated with a significant decrease in downstream expression levels (green downward triangles, Fig. 3b) in Pool B compared to Pool A (wildtype). The spike protein isoforms from Pool B showed lowered expression by at least five-fold and were associated with consecutive long-sequenced reads that contained a series of SNVs including at the position 23403 SNV, two A-to-G variants at position 21562 (1 nt before the AUG start site of S1 subunit) and position 22460 within the S1 subunit, as well as a G-to-U variant at position 25563 downstream of the S2 subunit in ORF 3 (Fig. 2, SG-Pool B asterisks, Fig. 3b, green triangles, and Fig. 4a, blue asterisks).

### Corroboration of sequence single nucleotide variant using ddPCR

In total, sixteen high quality SNVs were identified--in seven cases more than one SNV was found to be expressed at a particular site, suggesting that these may be more polymorphic regions (Fig. 4) and represent different the patients from each Pool. We validated the presence of three SNVs and their expression levels using ddPCR in comparison to the ATCC and Twist synthetic RNA controls and the SARS-CoV-2 Wuhan reference (MT008022) genome (Fig. 3-4a asterisks). Position 8782 (C>U) was confirmed as a wildtype “C” in Pool B and as expected, found to be the variant “U” in the ATCC control (Fig. 4b); resulting in a synonymous mutation. In Pool A, position 23403 (A>G) was confirmed in the spike S1 subunit of the ATCC control and matched the reference genome “A” nucleotide (Fig. 4c) whereas in Pool B had the “G” variant (Fig. 4c, bottom panel). The latter of these SNVs result in an amino acid change, D-to-G. ddPCR also revealed (and corroborated) the absolute levels of these isoforms in the pools. At position 14407 (nsp 12b), the “C” nucleotide was confirmed in the Twist RNA control, while in Patient 300 (part of Pool B), the “U” variant is observed. This latter variant results in an amino acid change P-to-L.

## DISCUSSION

Long-read sequencing identified a diverse transcriptional landscape in COVID-19 patients compared to what has been described previously. A wide range of transcripts, and isoform expression, was uncovered ranging 300-5000 nt in length, including detection of novel transcript isoforms and differentially expressed isoforms related to newly identified SNVs. The recombined architecture of SARS-CoV-2 was previously described by *Kim et al*., *2020* in Vero-infected cells using Nanopore and Nanoball sequencing, which allowed for the identification of discontinuous transcription events^8^. Similar to our observations of the ‘splicing’ of a shorter 65-70 nt 5’ UTR (instead of full-length 265 nt annotated region) with various downstream ORFs, advancing a ‘fusion hypothesis’, Kim et al., suggest that viral transcription in COVID-19 includes a negative-stranded intermediate form that facilitates fusion via the pausing and switching of RdRP machinery at the ∼10 nt transcription regulator sequence (TRS) body to TRS leader sites located at ORF junctions^9^. There are over 400 TRS described among different types of viruses, approximately four in coronaviruses^19^, where they can regulate viral transcript production^20^. In parallel, Solinska et al. (2011) noted that positive strand RNA viruses, including coronaviruses, use subgenomic RNAs as mRNAs, as observed in these long-sequence read data, whose additional properties could affect replication, transcription, gene rearrangements, and acquisition of non-self sequences^21^.

Many single nucleotide variants were identified through long reads, from which three were corroborated using ddPCR. Position 23403, GAU to GGU, results in an amino acid change aspartic acid (D) to glycine (G) in the spike protein (S:D614G). Interestingly, this glycine amino acid change is associated with increased infectivity, improving spike protein density, and facilitating entry into host ACE2 cells more effectively^22^. This variant was also described by Nextstrain analysis^23^ suggesting that patients in Pool B can be traced to an earlier occurrence (January 2020) than was first reported in the Washington State strain source. Interestingly, nine non-synonymous mutations in the spike protein have been of particular interest, having been characterized during the first spread of COVID-19 in Europe (Nextstrain). Our analysis, identified additional new spike variants in Pool B, including an A-to-G amino acid modifying at position 22460 (S:K300E), and one found in the spike S1 5’UTR, 1 nt upstream of spike start codon at position 21562 suggesting a possible regulatory role. Similarly, a structural role could be linked to the identified variant at position 241 in the 5’ UTR of Pool B, associated to a phylogenetic branch point by Nextstrain^23^. Overall, there have been reports of SARS-CoV-2 mutations thought to drive pathogenicity, several of which are found within the spike protein region with corresponding changes in the coding amino acid^22,24^. It will be of interest to further investigate the 3D protein structure in context to these amino acid changes.

Notably, the majority of SNVs were found to be A-to-G or C-to-U deaminase changes reminiscent of RNA editing events A-to-I ADAR and C-to-U APOBEC machinery. This pattern was observed when SARS-CoV-2 bronchoalveolar lavage fluid-based RNA sequence data was compared with those of other coronaviruses including MERS-CoV and SARS-CoV-1^25^. In their work, the authors proposed several mechanisms for SNV changes, including one where RNA editing occurs in coordination with negative strand transcription events^25^. Several variants we found showed partially expressed RNA editing, and several were located close to viral transcript rearrangement junctions associated to transcripts of variable gene expression. This points towards biological pathways operating in parallel to post-transcriptional modification events and secondary/tertiary RNA structures being critical for the recognition of specific consensus sites and possibly regulating the mechanisms by which viral transcript rearrangements and differential transcription events take place^26,27^. Distinct RNA structures and relatively high frequency of RNA viral recombination events in the family of Coronaviruses suggest a mechanistic regulation of viral genome organization rather than a selection of an evolutionary advantageous genotype^28^. Notably, distinct secondary structures are observed in coronaviruses, particularly the 5’ and 3’ UTRs^16,29^ which could additionally be involved in reassembly or fusion-like events. Spike protein variants are of particular interest, as they are distinct from other coronaviruses^8^ and vaccine and drug development have been based on these regions^30^.

Overall, the topography of the SARS-CoV-2 transcriptome is quite variable whereby the long-read method of sequencing and analysis give clearer perspectives on the distinct isoforms consisting of novel gene arrangements and SNVs, with varied expression and complexity, within and among the SARS-CoV-2 patient samples. These observations of diverse viral transcripts suggest that RNA editing, and recombination are relatively common occurrences and are likely part of viral biological mechanisms for transcript organization in patients positive for COVID-19^28^. Further investigation will be necessary to identify whether the abundance and reassortment of this viral transcription landscape varies and has patterns associated with severity and persistence of the viral infection, such as the control of infection in asymptomatic patients versus uncontrolled spread in patients with severe COVID-19-related pathology.

## Materials and Methods

### RNA extraction from nasopharyngeal swabs and Quality Control (QC)

The nasopharyngeal RNA swabs (in viral transport media, NP-VTM) from San Francisco Bay Area COVID-19 positive patient samples (diagnosed by Stanford Clinics) were stored in 1:4 Ratio of DNA/RNA shield to NP-VTM (total volume = 400uL) at −80°C prior to nucleic acid extraction representing one tenth of the volume of the original nasopharyngeal swab sample. The AllPrep DNA/RNA/Protein Mini Kit (Qiagen 80004) was used to extract total RNA in DEPC-treated water. The quality of the total RNA was tested using the Agilent Bioanalyzer 2100 (RNA 6000 Pico assay) noting that detection size range is 25 nt to10 kb, and the detection sensitivity range is 25-500 ng/ul. This concentration was also verified and validated using Qubit 2.0 Flourometer (RNA HS Assay Kit).

A total of 20 of the 24 samples showed relatively high quality of RNA when analyzed on the Bioanalyzer, whereby lower quality showed more degraded and fragmented RNA (Supplementary Fig. 1c and d). The Cycle threshold (Ct) was previously measured and compared to the RNA concentrations. Six distinct patient samples were selected to have the highest concentrations and quality and two sets of three samples of total real-world samples RNA were combined, Pool A (Patients 25, 125, 290) and Pool B (Patients 90, 260, 300) and concentrated in the SpeedVac for 15 min; with sample Ratios of Pool A (0.48, 0.32, 0.21) and Pool B (0.52, 0.12, 0.35), respectively.

### cDNA synthesis and amplification for long-read sequencing

A total of four RNA samples were prepared for cDNA synthesis. Two SARS-CoV-2 RNA synthetic controls were utilized and run in parallel with the Pools A and B for this work: 1) a Twist Bioscience SARS-CoV-2 (MT199235 – USA/CA9/2020) synthetic control containing six non-overlapping 5 kb fragments and 2) SARS-CoV-2 RNAs isolated from Vero E6 mammalian cells transfected with a 30 kb cloned isolate (MN985325.1 −2019-nCoV/USA-WA1/2020) supplied by ATCC (ATCC® VR-1986D™).

First strand cDNA synthesis was modified from: https://www.pacb.com/wp-content/uploads/Procedure-Checklist-Iso-Seq-Express-Template-Preparation-for-Sequel-and-Sequel-II-Systems.pdf and performed on two pooled samples, Pool A and Pool B of total RNA 228 ng and 165 ng, respectively. In parallel, ∼325 ng of ATCC and ∼380 ng of Twist SARS-CoV-2 controls were used. Universal Human Reference RNA (UHRR, Agilent) was spiked into each of the samples for a final RNA amount of 300 ng to balance the RNA concentrations in the pools. Total RNA was mixed with Reaction Mix 1 (9:1 of 12 uM NEB RT random 6N-S to 12 uM oligo-dT(20), 12uM dNTPs) with a final volume of 9 uL incubated 5 min at 70°C, cooled to 4°C, mixed with 10uL reaction mix 2 (containing NEBNext Single Cell RT Buffer and NEBNext Single Cell RT Enzyme Mix) incubated at 42°C for 75 min. The samples are cooled on ice prior to addition of Iso-Seq Express Template Switching Oligo and an additional 15 min incubation at 42°C for 15 min for a final volume of 20uL. The quality of the cDNA is assessed using Agilent Bioanalyzer 2100 (as above) and the COVID-19 positivity was corroborated by a new ddPCR method (described below). ProNex beads were used to clean up the RT-PCR product and eluted in 45.5uL EB buffer.

cDNA amplification was prepared with Reaction Mix 3: 50uL NEB Single Cell cDNA PCR Master Mix, 2uL NEBNext Single Cell cDNA PCR Primer, 2uL Iso-Seq Express cDNA PCR Primer and 0.5 NEBNext Cell Lysis Buffer. Cleaned first strand cDNA (eluted above) was combined with Mix 3 for a final volume of 100uL. PCR program was set up as follows: 1 cycle at 45 sec at 98°C, 20-25 cycles (10 sec 98°C, 15 sec 62°C, 3 min at 72°C), 1 cycle at 5 min at 72°C, hold at 4°C. Qubit 2.0 was used to determine that 25 cycles of PCR gave optimal concentrations to continue to next steps. All NEBNext reagents are from New England Biolabs (NEBNext Single Cell/Low Input cDNA Synthesis & Amplification Module (NEB #E6421S). 1x ProNex beads was used to purify the amplified cDNA and eluted to a final volume of 50uL in EB.

### Capture of SARS-CoV-2, Library preparation, Long-Read sequencing and Data Collection

500ng of cDNA per sample was input into the probe based capture reaction with the IDT COVID-19 biotinylated probes, capture protocol was modified from https://www.pacb.com/wp-content/uploads/Procedure-Checklist-%E2%80%93-cDNA-Capture-Using-IDT-xGen-Lockdown-Probes.pdf. The custom SARS-CoV-2 biotinylated probes consisted of 498 probes,120 nt in length and overlapped every 60 nt represented WuHan reference genome (MN908947.3). Post-captured and re-amplified, SMRTBell libraries were generated following the Iso-Seq Express protocol https://www.pacb.com/wp-content/uploads/Procedure-Checklist-Iso-Seq-Express-Template-Preparation-for-Sequel-and-Sequel-II-Systems.pdf. SMRTBell templates were bound to sequencing polymerase using the Sequel Binding and Internal Control Kit 3.0 (PN: 101-626-600). One Sequel 1M v3 LR SMRT Cell (20 hour movie) per library was sequenced on the PacBio Sequel platform using Sequel Sequencing Kit 3.0 (PN: 101-597-900).

### ddPCR Method to Detect SARS-CoV-2

The PCR primer pair and fluorescently labeled primer probes are listed in Supplementary Table 3.Thank-you to the Protein and Nucleic Acid Facility (PAN, Stanford School of Medicine) for probe and oligo synthesis. Digital Droplet PCR (ddPCR) to detect SARS-CoV-2 method was adapted from previous work in the Snyder Lab^31^ and developed to parallel qPCR Molecular Methods (Gold-Standard, used worldwide) with the additional advantage of multiple channel detection allowing for i*nvivo* SNV validation and quantification.

First strand cDNA synthesis for ddPCR was performed on Pool A, Pool B, and SARS-CoV-2 synthesized controls ATCC and Twist RNA samples. Mix 50 ng (range 1 ng-5 ug) of total RNA from each sample, 1 uL random hexamers (50uM, N8080127, Invitrogen), 1 uL of 10 uM dNTP mix and RNase free water for final volume of 12 uL and denature at 65°C for 5 min followed by immediate chill on ice. Briefly spin down and mix in 4uL 5X First-Strand Buffer (250 mM Tris-HCl, pH 8.3; 375 mM KCl; 15 mM MgCl_2_), 0.1 M DTT, 2 uL 0.1M DTT, and 1uL RNaseOUT™ (40 units/μL, #10777-019) and incubate at 25°C for 2 min. Add 1 ul of 200 U SuperScript™ II RT (#18064014, Invitrogen) for a final volume of 20 uL, and incubate at 25°C for 50 min. Inactivate the reaction by heating at 70°C for 15 min and chill on ice, with a final spin down to collect the cDNA sample. cDNA prepared for long-read sequencing can also be used. The annealing temperature was adjusted when gene specific reverse primers were used instead of random hexamers for this cDNA step.

For the digital droplet PCR step, prepare 20X PCR primer/probe stocks (for final concentration of 18 uM forward primer, 18 uM reverse primer, 5 uM FAM probe and 5 uM HEX or VIC probe. Prepare the ddPCR reaction by adding the reagents together: 10uL 2X Digital Super Mix containing polymerase enzyme, dNTPs and buffer (#110026868, BioRad), 1uL 20X PCR primer/probe stock, 1 uL cDNA from previous step (Pool A, Pool B and ATCC and Twist controls), and make to 20 uL with RNase-free water. The ddPCR droplets were prepared using the QX200 Droplet Maker Machine and Droplet Generation Oil (#10040718, BioRad). Thermal cycling protocol: 95°C for 10 min; 94°C for 30 seconds and 60°C for 1 min x 40 cycles (set to 50% ramp speed) and cooled prior to droplet measurement using the QuantaLife/BioRad QX200 Droplet Reader (Poisson distribution).

### Bioinformatic analysis and viewer

IGV_2.8.0 Browser^32^ was used to view and analyze the long-read sequences. StringTie_v2^33^ was used to identify and quantify (FPKM) long-read gene rearrangements (Supplementary Table 4). StringTie v2.1.4 usage included long reads processing (-L) which also enforces -s 1.5 -g 0 (default:false) and reference annotation (-G) used for guiding the assembly process (GTF/GFF3).

#### Long-Read analysis pipeline (LORE) development for SARS-CoV-2 virus transcriptome analysis

While StringTie and IGV analysis were efficiently identifying gene rearrangements in consecutive order, a subset of reads also contained novel sequences and gene rearrangements. To further characterize the transcriptome landscape of real-world COVID-19 positive samples, a custom bioinformatic pipeline was designed. The LORE Pipeline (https://github.com/jennlpt/LORE_longreadseqviral) is subdivided into three Parts (**Supplementary Fig. 3**): Part A-Identification and read-depth of SARS-CoV-2 transcripts and types of isoform rearrangement (including poly A), Part B-Identification and read-depth of novel reads (and repeats), and Part C-Identification and quantification of isoforms with SNV at ORF junction and associated expression of downstream isoforms.

#### <u>LORE Part A</u> (characterize unusual gene rearrangements and polyA tails)

A1) Select Identifiers from Reference genome (consecutive order based on reference genome). A2) Use Identifiers to-probe the transcripts (*.bam/*.sam output file). A3) Locate and quantify the number of transcripts that contain the sequence of identifiers

#### <u>LORE Part B</u> (Characterize novel reads and/or repeats)

B1) Identify novel reads as subsequences (not in SARS-CoV-2 reference genome). B2) Localization of subsequence within the long-read sequence reads (beginning of read up 30%, end of read greater >70% for reads greater than 3000 nt). B3) Identify, locate and quantify if subsequences are repeated. B4) Verification against NCBI database (https://blast.ncbi.nlm.nih.gov/). B5) Filter out non-viral related genes (see nRep below). B6) Optional: secondary structure of RNA using RNAfold (http://rna.tbi.univie.ac.at/cgi-bin/RNAWebSuite/RNAfold.cgi).

#### <u>LORE Part C</u> (SNVs and isoform relationship)

SNVs are identified using IGV_2.8.0 Browser^32^. Alignment replicated using bowtie2 (version 2-2.3.1) and SNVs called using samtools mpileup (samtools-1.9). SNVs with minimum depth coverage 25 and minimum unique read coverage of 3 were selected as high quality (as summarized in Supplementary Table 2). C1) Identify junction sites from the reference genome and categorize SNV locations within 10 nt from the junction. Discern novel-splice junctions^34^. C2) Calculate differential expression level downstream of isoforms containing SNVs relative to wildtype. RPKM = (CDS read count * 10^9^) / (CDS length * total mapped read count). FPKM or Fragment read is used when data is paired then only one of the mates is counted.

Additional sequences (nRep) that are associated with the long-read PacBio platform as adaptor sequence were identified using LORE (Part B). These nReps were located upstream and downstream of known genes in 25-28% of the samples including controls: namely nRep1 a ∼66 nt sequence “GGCAAUGAAGUCGCAGGGUUGUACUCUGCGUUGAUACCACUGCUUCCCUGUGGUUGUACGUCAAGG” comprised of 3 core elements (A-B-C) and the entire nRep1 segment is repeated (at least in part) up to 12 consecutive times and was up to 300 nt in length; nRep2 consists of a 25nt middle subsection of nRep1, **“**GUACUCUGCGUUGAUACCACUGCUU” (middle core element, B-B) repeated up to eight time; nRep3 is also a subsequence of nRep1 “GGCAAUGAAGUCGCAGGGUU” (first core element, A-A) repeated up to nine times and was mostly found upstream of the 5’ UTRs; and nRep4 is a combination of the core elements 2 and 1 (B-A), “GUACUCUGCGUUGAUACCACUGCUUCGGCAAUGAAGUCGCAGGGUU”. Overall, different mixtures of these cores are found upstream of the SARS-CoV-2 ORFs, in particular nsp 3. Interestingly, in four cases nRep1 sequences were located in the middle of SARS-CoV-2 ORFs (out of order) or with exogenous genes (mammalian RAB and SOD2) suggesting these exogenous adaptor primers form concatemers and can fuse with both viral and non-viral genes. These may also be artifacts of reverse transcriptase polymerase skipping to novel sequences, partly due to the predicted secondary structure of the repeat regions observed. This was also observed in other coronavirus examples, particularly with ORFs at the 5’ end of the genome, including a mouse coronavirus model that manifests viral hepatitis^15,35^. Consequently, these nReps are filtered from the raw data file in the pipeline.

## Acknowledgements

We thank Ting Hon, Jason Underwood, and Elizabeth Tseng of Pacific Biosciences for designing and performing the probe capture-based library protocol on the PacBio sequencing systems. We also would like to thank the Stanford Protein and Nucleic Acid (PAN) facility for probe and primer synthesis.

## Contributions

JLPT and SB performed the experiments. MKS contributed to sample preparation, QC and sample preparation. JLPT conceptualized, designed bioinformatic analysis tools and wrote the first draft of manuscript. All of the authors contributed to revising the manuscript.

## References

1. Cui, J., Li, F. & Shi, Z. L. Origin and evolution of pathogenic coronaviruses. Nat. Rev. Microbiol. 17, 181–192 (2019).

2. Cebey-López, M. & Salas, A. Recognizing the asymptomatic enemy. Lancet Infect. Dis. 3099, 20–21 (2020).

3. Hu, B., Guo, H., Zhou, P. & Shi, Z. L. Characteristics of SARS-CoV-2 and COVID-19. Nat. Rev. Microbiol. (2020) doi:10.1038/s41579-020-00459-7.

4. Walls, A. C. et al. Structure, Function, and Antigenicity of the SARS-Structure, Function, and Antigenicity of the SARS-CoV-2 Spike Glycoprotein. Cell 180, 281-292.e6 (2020).

5. Wan, Y., Shang, J., Graham, R., Baric, R. S. & Li, F. Receptor Recognition by the Novel Coronavirus from Wuhan: an Analysis Based on Decade-Long Structural Studies of SARS Coronavirus. J. Virol. 94, 1–9 (2020).

6. Shang, J et al. Structural basis of receptor recognition by SARS-CoV-2. 581, 221–224 (2020).

7. Lu, R. et al. Genomic characterization and epidemiology of 2019 novel coronavirus: implications for virus origins and receptor binding. Lancet 395, 565–574 (2020).

8. Chan, J.F. et al. Genomic characterization of the 2019 novel human-pathogenic coronavirus isolated from a patient with atypical pneumonia after visiting Wuhan. Emerg. Microbes Infect. 9, 540 (2020).

9. Andersen, K. G. et al. The proximal origin of SARS-CoV-2. Nat. Med. 26, 450–452 (2020).

10. Sola I., Almazán F., Zúñiga S., & Luis, E. Continuous and Discontinuous RNA Synthesis in Coronaviruses. Annu Rev Virol. 2, 265–288 (2015).

11. V’kovski, P., Kratzel, A., Steiner, S., Stalder, H. & Thiel, V. Coronavirus biology and replication: implications for SARS-CoV-2. Nat. Rev. Microbiol. (2020) doi:10.1038/s41579-020-00468-6.

12. Lauring, A.S. & Hodcroft, E. B. Genetic Variants of SARS-CoV-2 — What Do They Mean? JAMA Publ. online January 06, 2021. doi10.1001/jama.2020.27124 2, 6–8 (2021).

13. Mahase, E. Covid-19: What have we learnt about the new variant in the UK? 1–2 (2020) doi:10.1136/bmj.m4944.

14. Travers, K. J. et al. A flexible and efficient template format for circular consensus sequencing and SNP detection. Nucleic Acids Res. 38, p(2010).

15. Kim, D. et al. The Architecture of SARS-CoV-2 Transcriptome. Cell 181, 914-921.e10 (2020).

16. Yang, D. & Leibowitz, J. L. The structure and functions of coronavirus genomic 3’ and 5’ ends. Virus Res. 206, 120–133 (2015).

17. Lagarde, J. et al. High-throughput annotation of full-length long noncoding RNAs with capture long-read sequencing. Nat. Genet. 49, 1731–1740 (2017).

18. Dougherty, M. L. et al. Transcriptional fates of human-specific segmental duplications in brain. Genome Res. 28, 1566–1576 (2018).

19. Liu, X. et al. Human Virus Transcriptional Regulators. Cell 182, 24–37 (2020).

20. Bernard, H. U. Regulatory elements in the viral genome. Virology 445, 197–204 (2013).

21. Sztuba-Solińska, J., Stollar, V. & Bujarski, J.J. Subgenomic messenger RNAs: Mastering regulation of (+)-strand RNA virus life cycle. Virology 412, 245–255 (2011).

22. Zhang, L. et al. SARS-CoV-2 spike-protein D614G mutation increases virion spike density and infectivity. Nat. Commun. 11, 1–9 (2020).

23. Hadfield, J. et al. NextStrain: Real-time tracking of pathogen evolution. Bioinformatics 34, 4121–4123 (2018).

24. Yao, H. P. et al. Patient-Derived Mutations Impact Pathogenicity of SARS-CoV-2. SSRN Electron. J. (2020) doi:10.2139/ssrn.3578153.

25. Di Giorgio, S. et al. Evidence for host-dependent RNA editing in the transcriptome of SARS-CoV-2. Sci. Adv. 6, 1–9 (2020).

26. Li-Pook-Than, J. et al. Relationship between RNA splicing and exon editing near intron junctions in wheat mitochondria. Physiol. Plant. 129, 23–33 (2007).

27. Bishop, K. N., Holmes, R. K., Sheehy, A. M. & Malim, M. H. APOBEC-mediated editing of viral RNA. Science. 305, 645 (2004).

28. Simon-Loriere, E. & Holmes, E. C. Why do RNA viruses recombine? Nat. Rev. Microbiol. 9, 617–626 (2011).

29. Huston, N. C. et al. Comprehensive in-vivo secondary structure of the SARS-CoV-2 genome reveals novel regulatory motifs and mechanisms. Mol. Cell. 81, 1–15 (2021) doi:10.1016/j.molcel.2020.12.041.

30. Huang, Y., Yang, C., Xu, X. F., Xu, W. & Liu, S. W. Structural and functional properties of SARS-CoV-2 spike protein: potential antivirus drug development for COVID-19. Acta Pharmacol. Sin. 41, 1141–1149 (2020).

31. Chen, R. et al. Personal omics profiling reveals dynamic molecular and medical phenotypes. Cell 148, 1293–307 (2012).

32. Robinson, J. T. et al. Integrative Genome Viewer. Nat. Biotechnol. 29, 24–6 (2011).

33. Kovaka, S. et al. Transcriptome assembly from long-read RNA-seq alignments with StringTie2. bioRxiv 1–13 (2019) doi:10.1101/694554.

34. Zhang, Y., Liu, X., MacLeod, J. & Liu, J. Discerning novel splice junctions derived from RNA-seq alignment: A deep learning approach. BMC Genomics 19, 1–13 (2018).

35. Luytjes, W., Gerritsma, H. & Spaan, W. J. M. Replication of synthetic defective interfering RNAs derived from coronavirus mouse hepatitis virus-A59. Virology 216, 174–183 (1996).

