## Supplemental_Figs_Tables for "Long-read sequencing of SARS-CoV-2 reveals novel transcripts and a diverse complex transcriptome landscape"

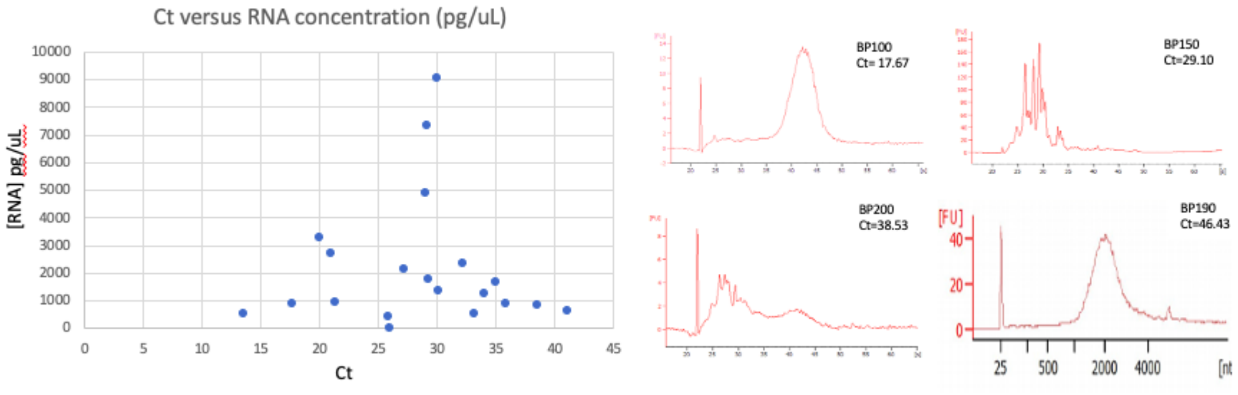

Supplemental Fig. 1: Cycle threshold (Ct) relative RNA concentration of COVID-19 positive samples. Six samples were selected for long-read sequences with relatively good quality (intact RNA) like BP-190 while those with more degraded appearance were not included in this work (BP150 and BP200).

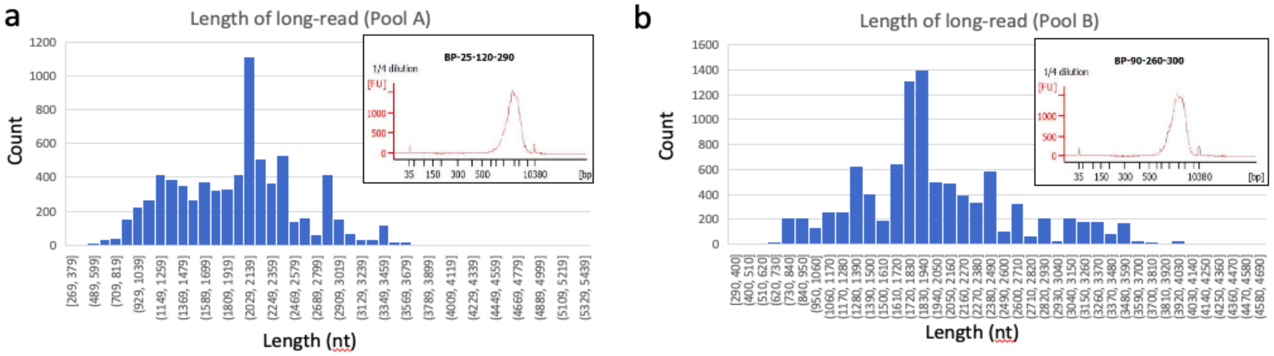

Supplemental Fig. 2. (a-b) Representation and length of Long-Read (Pacbio) transcripts from of COVID-19 positive patients from Pool A and Pool B, respectively. The corresponding Bioanalyzer graphs for Long-Read library data are shown (inset) showing the read-length distribution (x-axis), respectively.

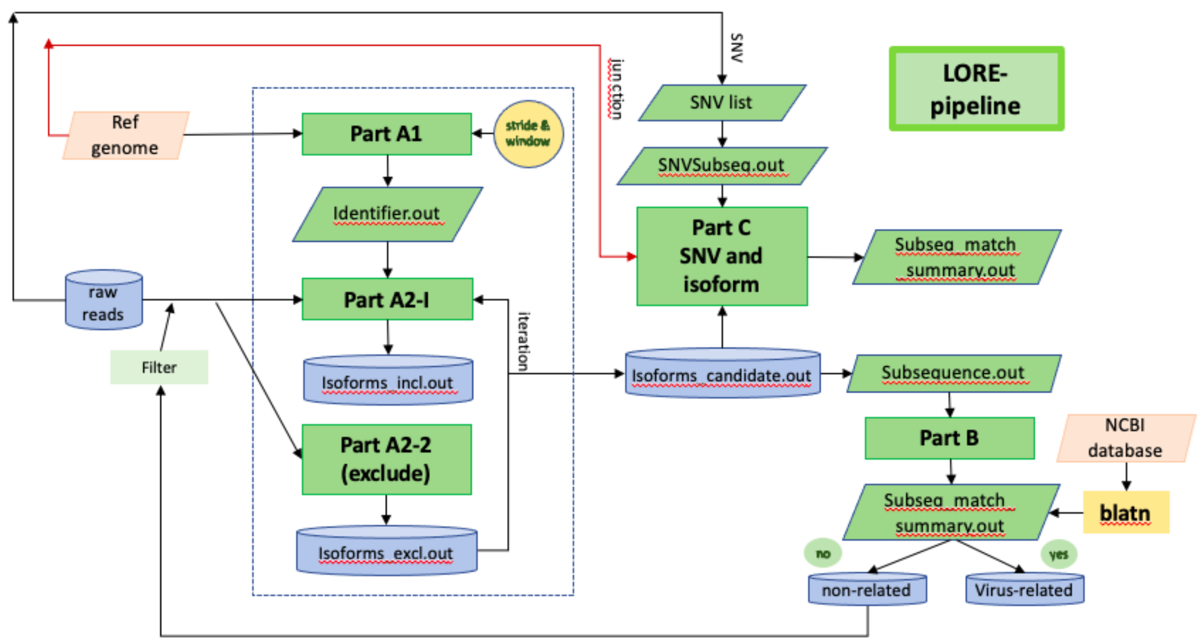

**Supplementary Fig. 3.** LORE pipeline. The three components of the LORE pipeline. Part A: Characterize gene arrangements. A1: Creates identifiers based on the Ref genome. In the first iteration stride and window length are based on the average read length (i.e. 1800 nt and 20 nt respectively) and the full length of the reference genome is used (~30 kb). The stride and window are adjusted dynamically as a function of the isoform candidate set that is produced in each reiteration for Part A2. Part B: Identifies non-reference sequences whereby output files are compared against available NCBI database. Part C: Identifies SNVs with high confidence (read depth >25, unique read >= 3). The reference genome is updated with the SNV and expression levels (read count) representing consecutive reads containing the SNVs are examined: a) close to ORF junction sites and b) expression levels relative to pooled samples. The raw reads are the long-read sequence (*.sam/*.bam files), reference genome GenBank: MT008022.1.

**Supplementary Table 1.** Primer and probes used for ddPCR.

| Gene | Label | Sequence 5'to3' |
|---|---|---|
| Envelope | WU_FRD_Epro | ACAGGTACGTTAATAGTTAATAGCGT |
| Envelope | WU_REV_Epro | ATATTGCAGCAGTACGCACACA |
| Envelope | WU_FAM_Epro | FAM-ACACTAGCCATCCTTACTGCGC-BHQ |
| Nsp4 | WU_FRD_8782 | ACTCGTGACATAGCATCTACAGA |
| Nsp4 | WU_REV_8782 | GCACGACAAAACCCACTTCT |
| Nsp4 | WU_FAM_8782-C | FAM-GGTTTAGCCAGCGTGGTG-BHQ |
| Nsp4 | WAS_HEX_8782-T | HEX-GGTTTAGTCAGCGTGGTG-BHQ |
| Spike | WU_FRD_23403 | ACTTCTAACCAGGTTGCTGTTC |
| Spike | WU_REV_23403 | ACCTGTAGAATAAACACGCCA |
| Spike | WU_FAM_23403-A | FAM-CAGGATGTTAACTGCACAGAAGTC-BHQ |
| Spike | DEU_HEX_23403-G | HEX-CAGGGTGTTAACTGCACAGAAGTC-BHQ |
| RdRp-orf12 | WU_FRD_14408 | GACAGATGCATTCTGCATT |
| RdRp-orf12 | WU_REV_14408 | ATCCTGATTATGTACAACACC |
| RdRp-orf12 | WU_FAM_14408-C | FAM-CAGTGTTCCCACCTACAAG-BHQ |
| RdRp-orf12 | WAS_FAM_14408-T | FAM-CAGTGTTCCCACTTACAAG- BHQ |

Legend: WU= Wuhan-Hu-1 (China) Reference Genome (MT008022.1), WAS= Washington-1 (USA) Reference Genome (MN985325.1), DEU= Deutche (Germany) Reference Genome (MT270101.1); FAM= 6-Carboxyfluorescein, HEX=Hexachlorofluorescein, BHQ=Black Hole Quencher

| Position | SNV | reference | % SNV | Change | Quality | Comment |
|---|---|---|---|---|---|---|
| 241 | 998 | | 100% | C>U | pass, low | Fig. 4 |
| 1049 | 1005 | | 100% | C>U | pass, high | Fig. 4 |
| 2721 | 390 | | 100% | C>U | pass, high | Fig. 4. junction |
| 3037 | 678 | | 100% | C>U | pass, high | Fig. 4 |
| 5956 | 119 | 188 | 39% | C>U | pass, low | poly A or U rich region |
| 5957 | 120 | 187 | 39% | A>U | pass, low | poly A or U rich region |
| 5988 | 44 | 141 | 24% | C>U | pass, low | throw out |
| 7395 | 16 | 37 | 30% | U>A | no pass | throw out |
| 8092 | 244 | 10 | 96% | C>U,A | pass, low | poly A or U rich region |
| 8782 | 48 | | 100% | C>U | pass, high | Fig. 4 |
| 11075 | 32 | 115 | 22% | U>C | pass, low | poly A or U rich region |
| 14286 | 263 | 415 | 39% | C>U | pass, low | poly A or U rich region |
| 14407 | 691 | | 100% | C>U | pass, low | Fig. 4 |
| 14771 | 146 | 409 | 26% | A>G | pass, low | throw out |
| 14867 | 261 | 145 | 64% | U>C | pass, low | throw out |
| 17747 | 540 | 19 | 97% | C>U | pass, high | Fig 4. partial edit candidate |
| 17858 | 1156 | 18 | 98% | A>G | pass, high | Fig 4. partial edit candidate |
| 18060 | 1061 | 44 | 96% | C>U | pass, high | Fig 4. partial edit candidate |
| 19687 | 659 | 715 | 48% | C>U | pass, high | Fig. 4 |
| 21562 | 628 | | 100% | A>G | pass, high | Fig. 4 |
| 21628 | 125 | 467 | 21% | U>C | no pass | throw out |
| 21698 | 81 | 146 | 36% | U>C | no pass | throw out |
| 22460 | 25 | 89 | 22% | A>G | pass, high | Fig. 4. (Pool B: Patient 260) |
| 23403 | 140 | | 100% | A>G | pass, high | Fig. 4. junction |
| 25563 | 38 | | 100% | G>A | pass, high | Fig. 4 |
| 26360 | 593 | 437 | 58% | U>C | pass, high | Fig. 4 (Pool A: Combo Patient 125, 290) |
| 26494 | 21 | 70 | 23% | U>G | no pass | throw out |
| 26497 | 8 | 20 | 29% | U>C | no pass | throw out |
| 28144 | 525 | 2 | 100% | U>C | pass, high | Fig. 4 |
| 29871 | 242 | 671 | 27% | A>G | pass, low | poly A or U rich region |

**Supplementary Table 2.** Summary of RNA Single Nucleotide variants (SNV) and candidate RNA-editing events. Parameters for SNV candidates include: minimum depth coverage 25, minimum unique read coverage 3 and distance of SNV from read end cut off = 10 nt. Low quality calls were allotted to the SNVs if found adjacent to a poly A or U rich region. Red triangle denotes shared SNVs with ATCC synthetic RNA control.

| POOL A | Posn 1 | Posn 2 | Isoform | reference_id NC_045512.2 | FPKM or cov | TPM |
|---|---|---|---|---|---|---|
| transcript | 1 | 29903 | transcript_id "STRG.1.1" | cov "64.927162" | FPKM "4669.746094" | TPM "103485.789062" |
| exon | 1 | 64 | transcript_id "STRG.1.1" | exon_number "1" | cov "58.180923" | |
| exon | 28255 | 29903 | transcript_id "STRG.1.1" | exon_number "2" | cov "65.217255" | |
| transcript | 1 | 29903 | transcript_id "STRG.1.2" | cov "41.534939" | FPKM "2987.311035" | TPM "66201.507812" |
| exon | 1 | 64 | transcript_id "STRG.1.2" | exon_number "1" | cov "2.529605" | |
| exon | 26468 | 29903 | transcript_id "STRG.1.2" | exon_number "2" | cov "42.261463" | |
| transcript | 1 | 29382 | transcript_id "STRG.1.3" | cov "27.923122" | FPKM "2008.310425" | TPM "44505.968750" |
| exon | 1 | 66 | transcript_id "STRG.1.3" | exon_number "1" | cov "16.111111" | |
| exon | 27385 | 29382 | transcript_id "STRG.1.3" | exon_number "2" | cov "28.287687" | |
| transcript | 1 | 29382 | transcript_id "STRG.1.4" | cov "25.036283" | FPKM "1800.680786" | TPM "39904.710938" |
| exon | 1 | 65 | transcript_id "STRG.1.4" | exon_number "1" | cov "8.456141" | |
| exon | 27884 | 29382 | transcript_id "STRG.1.4" | exon_number "2" | cov "25.755232" | |
| transcript | 1 | 265 | transcript_id "STRG.1.5" | reference_id "id-NC_045512.2:1 | cov "13.469845" | FPKM "968.789551" |
| exon | 1 | 265 | transcript_id "STRG.1.5" | exon_number "1" | reference_id "id-NC_045512.2:1..265" | cov "13.469845" |
| transcript | 266 | 29903 | transcript_id "STRG.1.6" | cov "1.616946" | FPKM "116.295349" | TPM "2577.209961" |
| exon | 266 | 20701 | transcript_id "STRG.1.6" | exon_number "1" | cov "1.592726" | |
| exon | 20860 | 29903 | transcript_id "STRG.1.6" | exon_number "2" | cov "15.738555" | |
| transcript | 29675 | 29903 | transcript_id "STRG.1.7" | reference_id "id-NC_045512.2:2 | cov "0.981875" | FPKM "70.619247" |
| exon | 29675 | 29903 | transcript_id "STRG.1.7" | exon_num | | |
| POOL B | | | | | | |
| transcript | 1 | 26468 | transcript_id "STRG.1.1" | cov "5520.067871" | FPKM "296157.343750" | TPM "855749.062500" |
| exon | 1 | 26468 | transcript_id "STRG.1.1" | exon_number "1" | cov "5520.067871" | |
| transcript | 1 | 29903 | transcript_id "STRG.1.2" | cov "324.295319" | FPKM "17398.779297" | TPM "50273.914062" |
| exon | 1 | 60 | transcript_id "STRG.1.2" | exon_number "1" | cov "16.986269" | |
| exon | 26494 | 29903 | transcript_id "STRG.1.2" | exon_number "2" | cov "329.702484" | |
| transcript | 29675 | 29903 | transcript_id "STRG.2.1" | :29675..29903", cov "0.726865" | FPKM "38.997059" | TPM "112.682312" |
| exon | 29675 | 29903 | transcript_id "STRG.2.1" | exon_number "1" | reference_id "id-NC_045512.2:29675..2 | cov "0.726865" |

**Supplementary Table 3.** Adapted Table representing StringTie Output files (version 2.1.4 was used). (sample usage: ./stringtie -L -G reference.gff.txt -o *output_file.gtf --viral *sample_data.mapped.bam).
